# Tissue Specific Requirement for the Inv(16) Oncogene *CBFB::MYH11* in Acute Myeloid Leukemia

**DOI:** 10.1101/2024.10.06.616896

**Authors:** Sipra Panda, Yiqian Wang, Venkatasai Rahul Dogiparthi, Michelle Becker, Arjun Dhir, Calvin Lam, Cecilia Rivas, Lemlem Alemu, Lisa Garrett, Peng Xiao, Samantha A. Swenson, Kyle J. Hewitt, R. Katherine Hyde

**Affiliations:** Department of Biochemistry and Molecular Biology, University of Nebraska Medical Center, Omaha, NE; Fred and Pamela Buffett Cancer Center, University of Nebraska Medical Center, Omaha, NE; Department of Genetics, Cell Biology and Anatomy, University of Nebraska Medical Center, Omaha, NE; Transgenic Mouse Core, National Human Genome Research Institute, NIH, Bethesda, MD; Translational and Functional Genomics Branch, National Human Genome Research Institute, NIH, Bethesda, MD

## Abstract

Inversion of chromosome 16 [inv(16)] generates the fusion gene *CBFB::MYH11 (CM)* and is one of the most common chromosomal rearrangements in Acute Myeloid Leukemia (AML). Expression of *CM* is required for leukemia initiation. Patients with inv(16) at diagnosis invariably have the rearrangement at relapse, leading to the assumption that *CM* is also required after leukemic transformation. However, a role for *CM* in leukemia maintenance has yet to be shown experimentally. To address this, we used an inducible *CM* knockdown (KD) mouse model and found that decreased *CM* eliminated leukemia cells from the peripheral blood and spleen, but not the bone marrow, despite all populations exhibiting significantly decreased *CM* mRNA and protein. The surviving *CM* KD cells in the bone marrow showed decreased apoptosis and proliferation, and increased expression of autophagy related genes. Surprisingly, with prolonged KD of *CM*, ∼40% of mice re-established disease despite maintaining decreased *CM*. Our work indicates that *CM* is required for leukemia survival in the spleen and peripheral blood, but in the bone marrow *CM* KD leukemia cells can survive and re-establish disease independent of the fusion protein. These findings imply that targeting *CM* alone has potential to reduce leukemic burden but not cure the disease.

## Introduction

Acute Myeloid Leukemia (AML) is a heterogeneous clonal malignancy arising from the hematopoietic stem and progenitor cells of the bone marrow, as the result of a block in differentiation that gives rise to a more than 20% increase in immature blast cells, reduced mature blood cell production and ultimately BM failure (1–3). AML develops as a neoplastic disease, often with specific chromosomal rearrangements, including the inversion of chromosome 16[inv(16)] (4, 5). These rearrangements are considered the initiating mutation, and with the acquisition of additional cooperating mutations, these abnormal myeloid cells can transform into a full-blown, frank leukemia (6–8). Inv(16) accounts for ∼10% of all AMLs, making it one of the most common chromosomal abnormalities in AML (9–12). At the molecular level, Inv(16) generates a fusion between the genes encoding the transcription factor Core Binding Factor β (*CBFB*) and the contractile protein Smooth Muscle Myosin Heavy Chain (*MYH11*). The resulting oncogene, *CBFB::MYH11 (CM),* encodes the protein CBFβ::SMMHC (13–15). With current treatment protocols, Inv(16) AML has a favorable prognosis due to high rates of initial response to chemotherapy. However, ∼40% of patients will eventually relapse, at which point there are limited curative options (12, 16, 17).

Because *CM* is the initiating mutation in this subtype of AML and is expressed in all leukemia cells but not in healthy cells, it is an ideal drug target. However, the role and requirement for *CM* after malignant transformation is poorly understood. In Chronic Myeloid Leukemia (CML), another fusion protein driven leukemia, expression of BCR::ABL initiates the disease, but once established, BCR::ABL activity is largely dispensable for maintenance of the leukemia stem cell population. Currently, it is not known if this is a unique feature of BCR::ABL and CML, or if the existence of a population of cells that no longer require the activity of the initiating mutation is a more common phenomenon. Resolving this issue is critical to the development and eventual clinical use of inhibitors targeting other leukemia fusion proteins like CBFβ::SMMHC in AML.

In this study, we investigate the role of *CM* in primary mouse leukemia cells after leukemia initiation. We found that decreased *CM* expression eliminates leukemia from the peripheral blood and spleen, but not the bone marrow, indicating that loss of CBFβ::SMMHC activity is sufficient to control, but not cure the disease.

## Methods

### Mice

All animal experiments were approved by the University of Nebraska Medical Center IACUC in accordance with NIH guidelines. 4-6-week-old mice of both sexes were used.

#### Mx1-Cre, Cbfb^+/56M^ mice

To generate *CM* expressing leukemia cells, we used conditional knock-in mice expressing *Cbfb::MYH11* from the endogenous promoter, (*Cbfb^+/56M^*) paired with the inducible *Mx1-Cre* transgene: *Mx1-Cre, Cbfb^+/56M^* mice (18). To induce leukemia, *CM* mice were treated with 250µg Polyinosine-polycytidylic acid (pI:pC) 3 times a week, for 3 rounds of treatment.

#### _Cbfb_+/flMYH11 _mice_

To generate knock-in mice with the *flMYH11* allele, *LoxP* sites were added by site directed mutagenesis (QuikChange, Agilent) flanking the *Cbfb Exon 5::MYH11* cassette in the *pPNT* vector. Mouse embryonic stem (ES) cells were electroporated, selected, screened for homologous recombination and injected into blastocysts, as described previously (19, 20). To confirm the *Cbfb^flMYH11^* allele functions as predicted, chimeric mice were bred to mice expressing *Cre* from the *β-Actin* promoter (*ActB-Cre^+^*), purchased from Jackson Laboratory (Bar Harbor, ME, USA) and embryos harvested at embryonic day 12.5 (21). To induce leukemia, chimeric *Cbfb^+/flMYH11^*mice were treated with a single dose of N-ethyl-N-nitrosourea (ENU, 100mg/kg) and monitored for leukemia development, as described previously (22–24).

#### Transplantation of leukemia cells

*CM* expressing leukemia cells were transplanted into sub-lethally irradiated congenic recipient mice by retro-orbital injection, as described previously (25, 26).

### Viral Transduction

To excise the *Cbfb ^flMYH11^* allele, lentiviral plasmids co-expressing *Cre Recombinase* and either GFP or DsRED were used. To knockdown *Cbfb::MYH11*, a doxycycline (Dox) inducible lentiviral plasmid expressing GFP and a shRNA against *MYH11* (*shMYH11*) was kindly provided by Dr. J. Martens, Nijmegen, NL (27). Virus was produced in 293KEK cells and used to transduce mouse leukemia cells, as previously described (25, 26, 28). Transduced cells were sorted for GFP or dsRED using Aria II Cell Sorter (BD Biosciences, San Diego, CA, USA) 48hrs post transduction and transplanted into congenic mice as above. To induce *shMYH11* expression, mice were given 5% sucrose supplemented drinking water with or without (CTRL) Dox at 2mg/ml. Mice were sacrificed on the indicated day of treatment and tissue collected.

### qRT-PCR Analysis

RNA was extracted using Trizol (Thermo Scientific, Waltham, MA, USA) as described previously (25, 26). cDNA was synthesized and amplified using Luna RT Supermix and Luna Universal qPCR kit (New England Biolabs, Ipswich, MA, USA) per manufacturer’s protocol. Primer sequences available upon request.

### Western Blot Analysis

Cell lysates were prepared, subjected to SDS-PAGE and transferred to PVDF, as described previously (25, 26). Membranes were blocked in 5% milk in TBST and then probed using primary antibodies and then with secondary HRP-linked antibodies as indicated in (Supplementary Table 1). Protein bands were visualized with SuperSignal West Pico PLUS (Thermo Scientific). Quantification was performed using ImageJ software.

### Flow Cytometry Analysis

Cells were stained with fluorophore coupled antibodies against B220 (RA3-6B2), CD3 (17A2), Mac-1/CD11b (M1/70), GR-1 (RM4-5), CKIT (53-6.7), Csf2rb/CD131 (JORO50), IL1RL1 (U29-93), CD41 (MWReg30), CD123 (5B11) (BD Bioscience) and analyzed by flow cytometry (BD, LSR II). Apoptosis was assayed using Annexin V (BD). Proliferation was determined using the BrdU Flow Kit (BD). Mice were injected with BrdU at dose of 200mg/kg body weight. After 1 hr, mice were sacrificed and tissue harvested. All flow cytometry data was analyzed using FlowJo 10.0 (BD, Ashland, OR, USA).

### Colony Assays

Equal numbers of cells were suspended in Methocult (m3434, StemCell Technologies, Vancouver, BC) in triplicate and plated in 35mm dishes, according to manufacturer’s protocol. Colonies were counted after 14 days.

### Whole Transcriptome sequencing

Total RNA was extracted as above. RNA was quantified using Qubit 2.0 Fluorometer (Thermo Scientific) and integrity was checked using Agilent TapeStation 4200 (Agilent Technologies, Palo Alto, CA, USA). Library preparation and sequencing was carried out by Azenta Life Sciences (Burlington, MA, USA). Further details about analysis provided in (Supplementary Methods).

### Single cell RNA sequencing

Leukemia cells were enriched using the MagniSort Mouse Hematopoietic Lineage Depletion kit (Invitrogen) then single cell RNA sequencing (scRNA-Seq) was performed using the Seq-well, nanowell-based high throughput platform, as described previously (29, 30). Details provided in the (Supplementary Methods).

### Statistical Analysis

Statistical significance was determined using either Student’s T-Test or ANOVA, as appropriate (GraphPad Prism, version 10, Dotmatics, Boston, MA, USA) with p-values ≤ 0.05 deemed significant. Sample size given in the figure legends.

## Results

### Knockdown of *Cbfb::MYH11* (*CM*) decreases leukemic burden in the spleen and peripheral blood, but not the bone marrow

To test the role of *Cbfb::MYH11 (CM)* in leukemia maintenance *in vitro*, we used leukemia cells from *Cbfb^+/56M^*, *Mx1-Cre^+^* mice which conditionally express *CM* from the endogenous *Cbfb* promoter and develop leukemia that faithfully recapitulates human Inv(16) AML (18). We collected and transduced these leukemia cells with a doxycycline (Dox) inducible shRNA against *MYH11* (*shMYH11*) which also constitutively expresses GFP (27). We sorted GFP^+^ cells and cultured them with or without Dox for 48 hours (hrs). As expected, we observed that Dox induced *CM* KD resulted in a significant decrease of *CM* expression at both the mRNA and protein levels (Fig. 1A-B). We also found that *CM* KD induced increased apoptosis and decreased colony formation (Fig. 1C, D). These findings indicate that *CM* is required for the growth and survival of frank leukemia cells *in vitro*.

**Figure 1:**
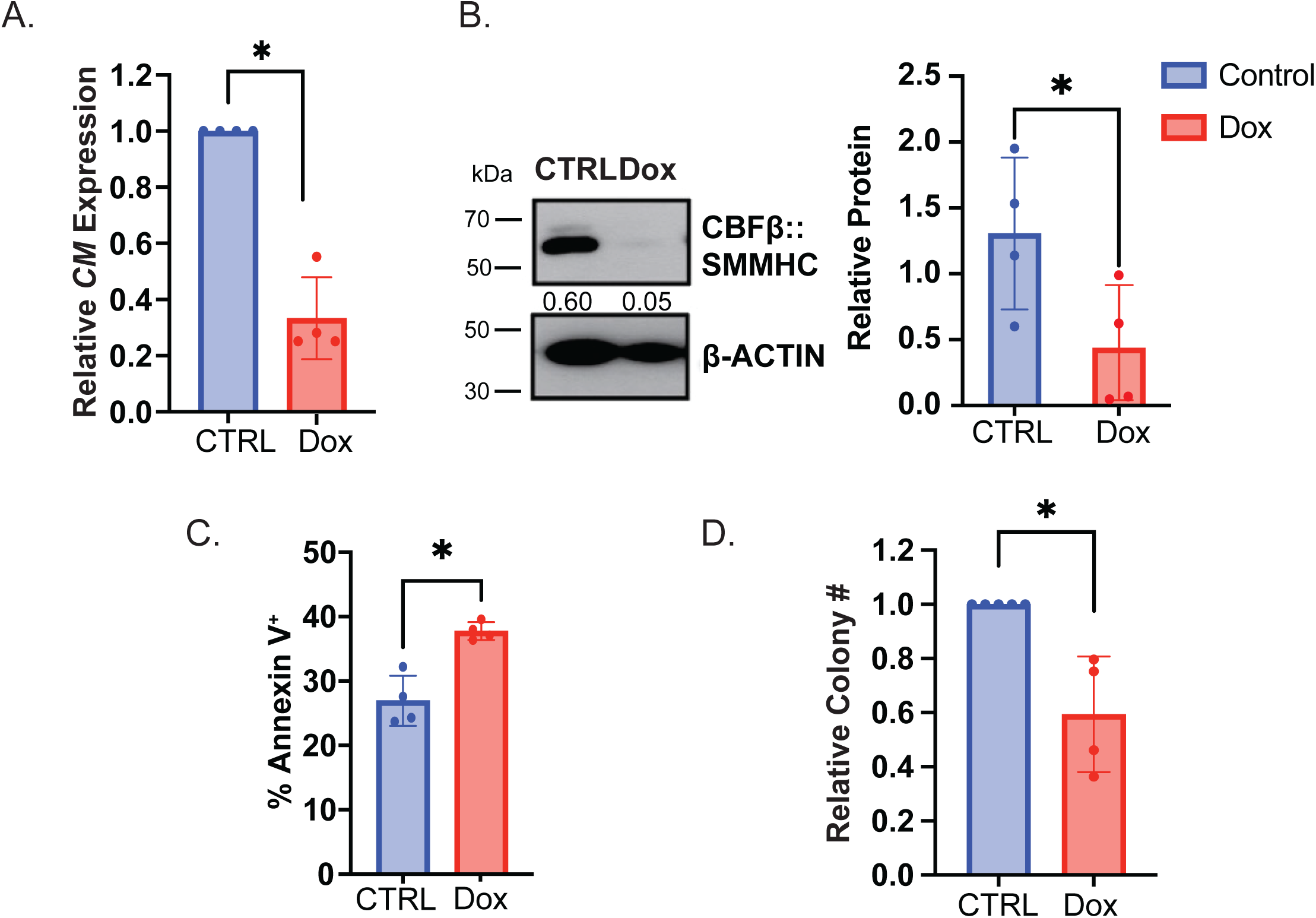
Inducible knockdown of *CM* causes apoptosis and impairs colony formation *in vitro*. **A. Bar** graph showing quantitative reverse transcription PCR (qRT-PCR) of *CBFB::MYH11* in *Cbfb^+/56M^* leukemia cells transduced with an inducible shRNA against *MYH11* (shMYH11) treated *in vitro* with either control (CTRL) or doxycycline (Dox) for 48 hrs **B.** Representative western blot (left) and bar graph of relative CBFβ::SMMHC protein levels (right) in *shMYH11* transduced leukemia cells treated as in A. **C**. Bar graph showing the percentage (%) of *shMYH11* transduced leukemia cells treated with control or Dox that are positive for Annexin V. **D.** Bar graph showing relative colony numbers generated by cells treated as in A. N ≥ 3 biological and technical replicates, * = p ≤ 0.05 compared to CTRL.

To determine the effect of decreased *CM* expression *in vivo*, we transplanted *shMYH11* transduced cells into congenic mice and allowed them to engraft. Once mice showed at least 20% GFP^+^ cells in the peripheral blood (Supplementary Fig. S1A), we induced *CM* KD and determined the percentage of GFP^+^ leukemia cells after 4, 7, 14, 21, and 28 days. We observed that *CM* KD induced a significant decrease in GFP^+^ leukemia cells in the peripheral blood and spleen starting at day 7, with near complete elimination by day 14 (Fig. 2A, B). In addition, we observed a notable decrease in spleen size with *CM* KD (Supplementary Fig. S1B, C). In the bone marrow, there was a significant decrease in leukemia cells starting at day 7 of Dox treatment. However, leukemia cells continued to make up 20-30% of the bone marrow through day 28 (Fig. 2C). To determine if the persistence of leukemia cells in the bone marrow was due to a failure of Dox to penetrate the tissue, we sorted GFP^+^ leukemia cells from the bone marrow of control and Dox treated mice at days 7 and 21. Quantitative real time PCR (qRT-PCR) and western blot analysis of bone marrow GFP^+^ cells showed significant *CM* KD at both time points (Fig. 2D-F).

**Figure 2:**
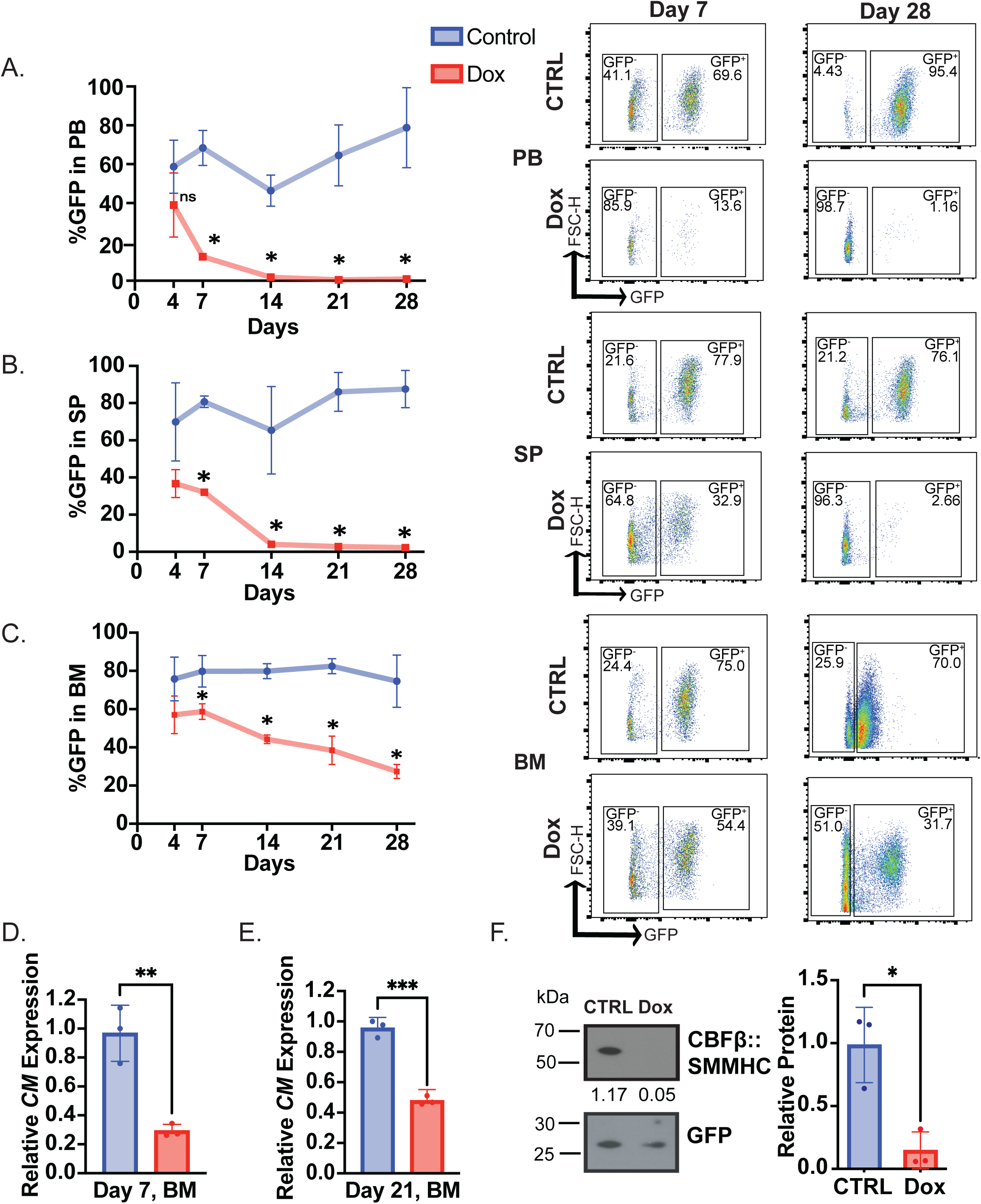
*CM* knockdown decreases leukemic burden *in vivo.* **A.** Line graph showing % GFP^+^ *shMYH11* transduced leukemia cells (left) and representative flow cytometry plots (right) in the peripheral blood (PB), **B.** spleen (SP) and **C.** bone marrow (BM) of mice treated with Dox or CTRL for 7, 14, 21, or 28 days. **D.** Bar graph showing relative expression of *CM* in sorted GFP^+^ leukemia cells from the BM of mice treated with Dox or CTRL for 7 and **E.** 21 days. **F.** Representative western blot (left) and bar graph (right) of relative CBFβ::SMMHC protein levels in sorted GFP^+^ cells from the BM of mice treated with either CTRL or Dox for 7 days. N ≥ 3 biological replicates, ns = not significant, * = p ≤ 0.05.

To confirm that the loss of leukemia cells in the Dox treated mice is due to decreased *CM* expression and not the effect of Dox, we used *CM* expressing leukemia cells transduced with a non-targeted shRNA construct. We observed no difference in the percentage of GFP^+^ leukemia cells between Dox and control treated mice (Supplementary Fig. S1D, E). These findings indicate that cells with reduced *CM* can survive in bone marrow, but not the peripheral blood or spleen.

### Deletion of *Cbfb::MYH11* impairs the growth and survival, but does not eliminate colony formation, of leukemia cells *in vitro*

To validate our finding that a population of leukemia cells can survive without the fusion gene, we generated a new knock-in mouse model that allows deletion of the *Cbfb::MYH11* fusion gene by Cre Recombinase. We used homologous recombination to knock-in the *Cbfb exon5::MYH11* fusion flanked by *LoxP* sites into the endogenous *Cbfb* locus (*Cbfb^+/flMYH11^*) (Fig. 3A). We identified embryonic stem cell (ES) clones with the knock-in allele by southern blot and confirmed CBFβ::SMMHC expression by western blot (Supplementary Fig. S2A, B). Three independent *Cbfb^+/ floxMYH11^* ES clones were used to generate chimeric mice. To test that the *Cbfb ^flMYH11^* allele can be fully excised, chimeric mice were crossed with mice expressing *Cre Recombinase* from the *Actin B* promoter (*ActB Cre*), which should allow for excision of the fusion gene in all cells of an embryo. We harvested embryos at day 12.5 of development. In *Cbfb^+/ flMYH11^* embryos lacking the *ActB-Cre* transgene, we observed central nervous system hemorrhaging, indicating expression of *Cbfb::MYH11*, as in previous knock-in models (Fig. 3B) (19). Importantly, this phenotype was not observed in *Cbfb^+/flMYH11^*; *ActB Cre^+^*, indicating excision of the fusion gene in the presence of Cre Recombinase. PCR using primers specific for the unexcised and excised alleles confirmed that the *Cbfb ^flMYH11^* was efficiently excised in *Cbfb^+/flMYH11^*; *ActB-Cre^+^* mice. To generate leukemia cells expressing the *Cbfb ^flMYH11^* allele, chimeric mice were treated with N-ethyl-N-nitrosourea and monitored for disease onset. Once leukemia was detected, cells were collected and transplanted into congenic recipient mice and monitored for survival. Recipient mice transplanted with leukemia cells from 3 different ES cell clones developed disease with a similar immunophenotype and histological appearance as previous *CM* knock-in mouse models (Fig. 3C, D; Supplementary Fig. S2C, D) (18, 31). These results indicate that *Cbfb^+/flMYH11^* mice develop a frank leukemia with a similar phenotype to previous models and that the fusion gene can be deleted by Cre Recombinase.

**Figure 3:**
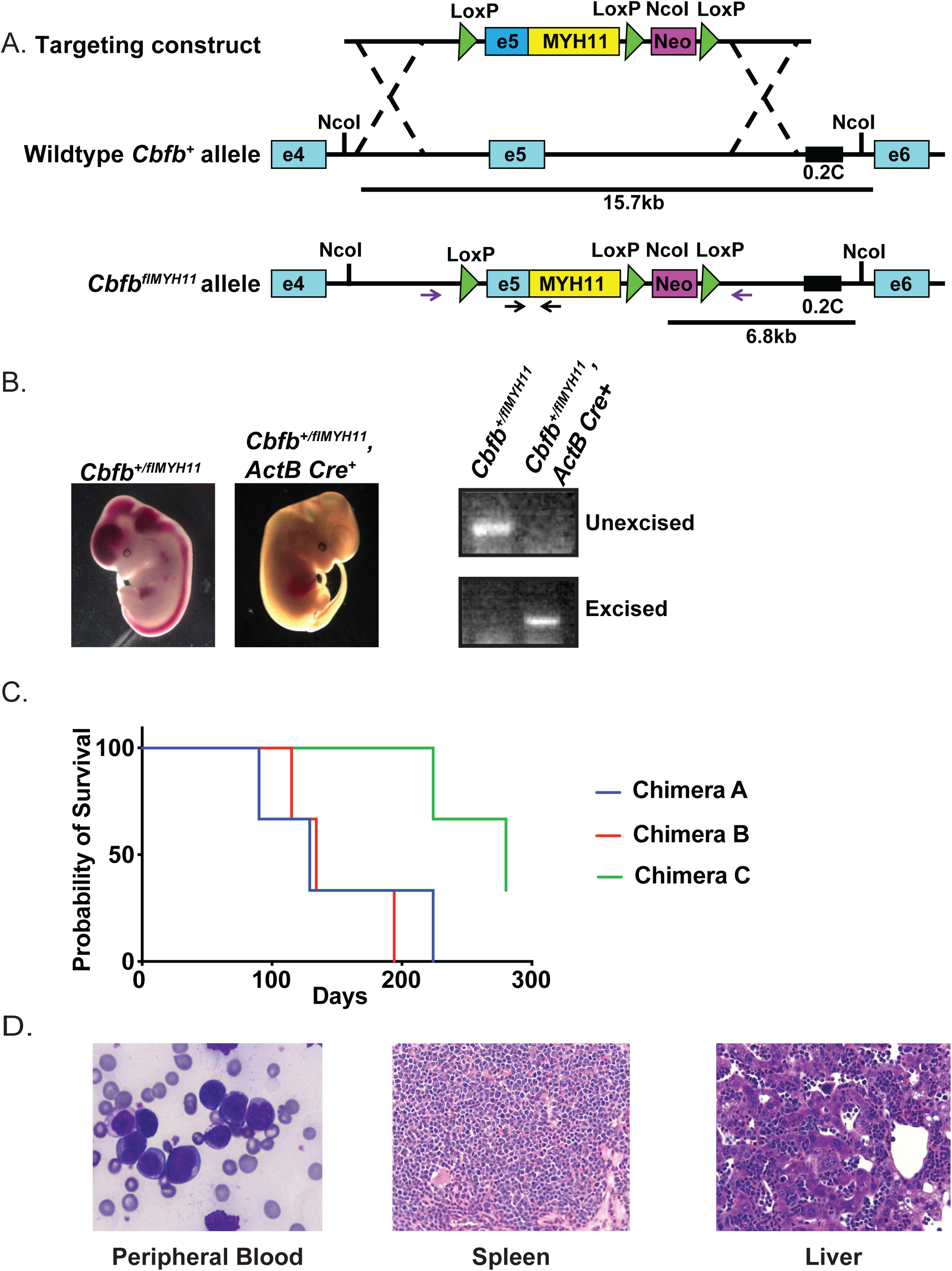
Generation of flox*Cbfb::MYH11* mice. **A.** Schematic representation of the embryonic stem (ES) cell targeting strategy. The probe (0.2C) and restriction enzyme sites (NcoI) used for screening are indicated. The black arrows represent PCR primers for the excised allele, and the purple arrows represent primers for the unexcised allele. **B.** Pictures of representative embryos harvested at embryonic day 12.5 generated by crossing *Cbfb^+/floxMYH11^* chimeras with transgenic mice expressing *Cre recombinase* (Cre) from the *β–Actin* promoter (*ActB Cre*). The right panel shows a representative PCR for the excised and unexcised *Cbfb^+/floxMYH11^* allele in DNA from embryos of the indicated genotype. **C.** Kaplan-Meyer survival curve of recipient mice transplanted with *Cbfb^+/flMYH11^* leukemic cells from three different chimeric donor mice. **D.** Representative Wright-Giemsa-stained peripheral blood smear and hematoxylin & eosin-stained spleen and liver section from a leukemic *Cbfb^+/flMYH11^* recipient mouse.

To test the effect of *CM* deletion *in vitro*, we transduced *Cbfb^+/flMYH11^* leukemia cells with a lentivirus expressing *Cre* and GFP from an IRES (Fig. 4A). Transduced cells were sorted, and *CM* excision was confirmed by qPCR and western blot (Fig. 4B-C). We found that *CM* deletion caused a statistically significant increase in Annexin V, indicating induction of apoptosis (Fig. 4D). To test the effect of *CM* deletion on colony forming ability, we plated equal numbers of control and *Cre* transduced *Cbfb^+/flMYH11^* leukemia cells in methylcellulose. After 14 days, colonies were counted, resuspended as single cells, and equal numbers of cells replated in methylcellulose. At both the 1st and 2^nd^ plating, we found that *Cbfb^+/flMYH11^* cells transduced with *Cre* generated significantly fewer colonies (Fig. 4E). To determine if the colonies that were able to grow from *Cre*-infected cells had successfully deleted *CM*, individual colonies were picked and qPCR for the *Cbfb^flMYH11^* allele was performed. We found that most colonies retained the unexcised *CM* allele. However, 5-20% of the colonies showed successful deletion of *CM* (Supplementary Fig. S3A, B). These findings are consistent with our *in vivo* knockdown data in which the majority of leukemia cells require continued expression of the fusion gene after leukemia initiation, but that there is a population of cells that are able to survive without *CM*.

**Figure 4.**
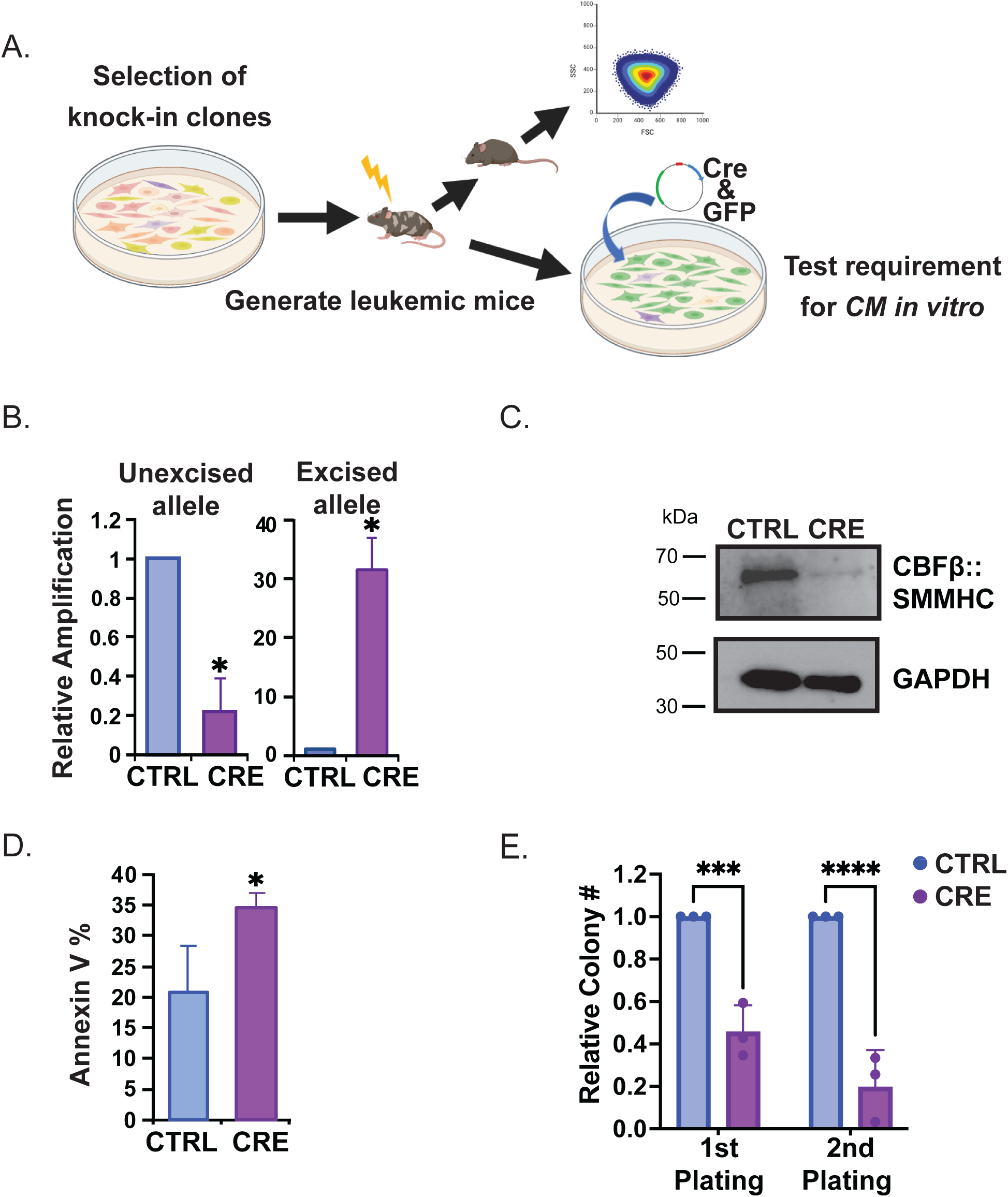
Deletion of *Cbfb::MYH11* induces apoptosis and reduces colony formation *in vitro*. **A.** Schematic representation of the experimental design to test the effect of *Cbfb::MYH11* deletion *in vitro*. **B.** Bar graph of qRT-PCR of the unexcised or excised *Cbfb^+/floxMYH11^* alleles in leukemia cells transduced with a lentivirus expressing Cre Recombinase (Cre) or Control (CTRL) and sorted for GFP. **C.** Representative western blot of CBFβ::SMMHC in *Cbfb^+/flMYH11^* leukemia cells infected with CTRL or Cre virus. **D.** Bar graph of % Annexin V in CTRL or Cre transduced *Cbfb^flMYH11^* leukemia cells. **E.** Bar graph showing the relative number of colonies from equal numbers of CTRL or Cre transduced *Cbfb^+/flMYH11^* leukemia cells. Colonies were scored on day 14. N = 3 biological and technical replicates, * = p ≤ 0.05, *** = p ≤ 0.001 as compared to CTRL.

### KD of *CM* alters apoptosis and proliferation

To investigate potential mechanisms for the persistence of *CM* KD cells in the bone marrow, we analyzed apoptosis and proliferation in the leukemic cells of Dox and control treated mice at Day 7 and Day 21. At Day 7, we found that Dox, but not control, treated mice had a significant increase in apoptosis in the spleen, but not the bone marrow (Fig. 5A left). However, at day 21, a time point in which control mice have high leukemic burden, we found high levels of apoptotic leukemia cells in the bone marrow of control treated mice, but not Dox treated mice (Fig. 5A right). To determine if the failure to induce apoptosis in the bone marrow with *CM* KD is related to the levels of anti-apoptotic proteins, we sorted leukemia cells from the spleen and bone marrow of control and Dox treated mice on day 7 and performed western blots. We found that KD of *CM* induced decreased BCL-2 in the spleen but caused no change in MCL-1. In leukemia cells from the bone marrow of both control and Dox treated mice, BCL-2 was barely detectable. In contrast, MCL-1 was higher in leukemia cells from the bone marrow of both control and Dox treated mice, as compared to cells from the spleen (Fig. 5B, C). These findings imply that MCL-1, but not BCL-2, may contribute to the survival of *CM* KD cells in the bone marrow.

**Figure 5:**
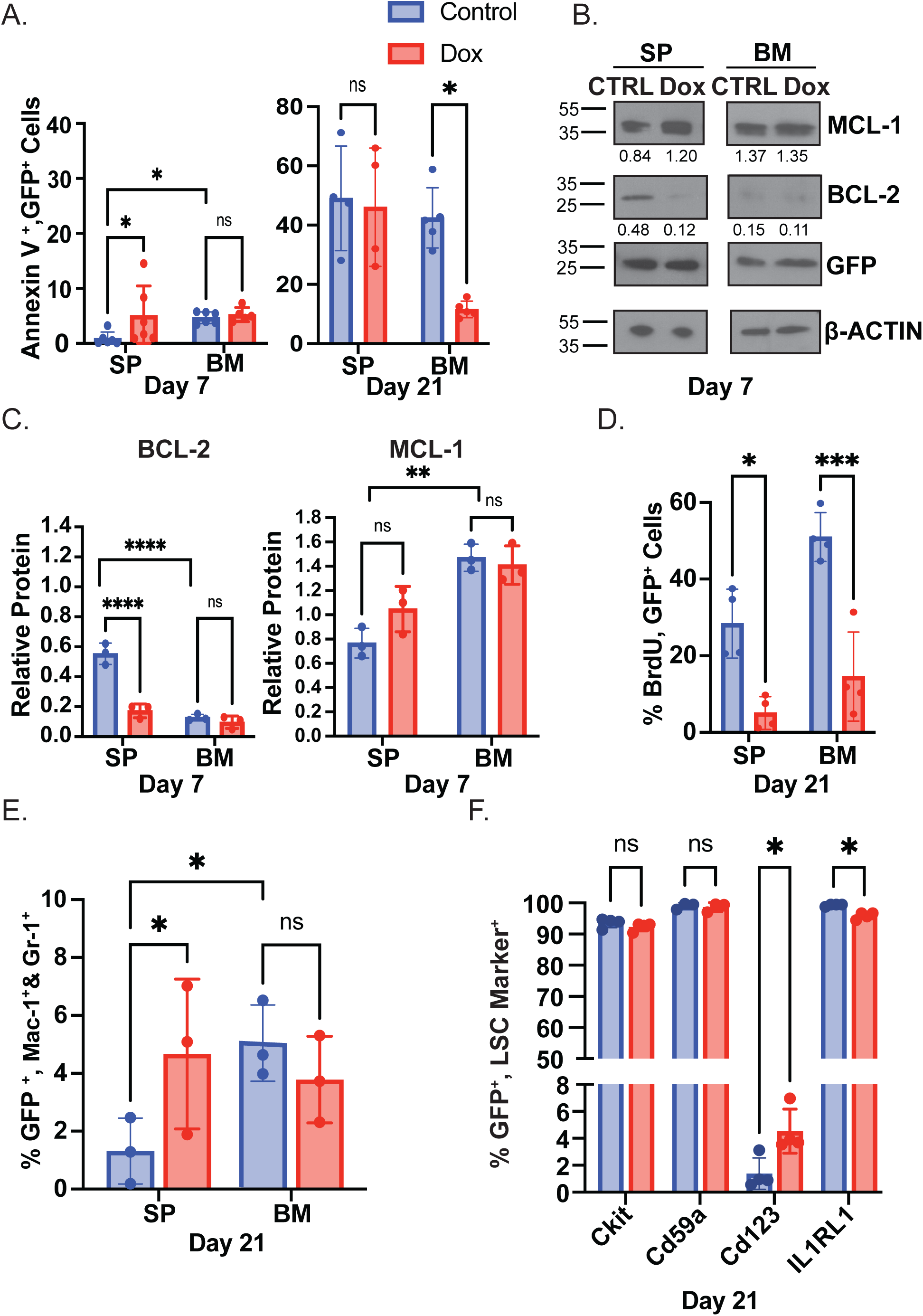
*CM* Knockdown induces apoptosis in leukemia cells in the spleen, but not the bone marrow. **A.** Bar graph showing the % Annexin V^+^, GFP^+^ cells from the SP and BM of mice treated with CTRL or Dox for 7 (left) or 21 days (right). **B.** Representative western blot and **C.** Bar graph of quantification for the indicated proteins from leukemia cells sorted for GFP from the SP and BM after 7 days of treatment with CTRL or Dox **D.** Bar graph of the % BrdU^+^, GFP^+^ leukemia cells harvested from the SP and BM of mice after 21 days of treatment with CTRL or Dox. **E.** Bar graph showing % GFP^+^ cells expressing the mature myeloid markers Mac-1 and Gr-1 from the SP and BM of mice treated as in D. **F.** Bar graph showing the % GFP^+^ cells expressing the indicated leukemia stem cell markers in the BM from of mice treated as in D. N ≥ 3 biological replicates, ns = not significant, * = p ≤ 0.05, ** = p ≤ 0.01, *** = p ≤ 0.001 as compared to CTRL.

To determine if a difference in proliferation could also contribute to the persistence of *CM* KD cells in the bone marrow, we used BrdU incorporation. Leukemia cells from both the spleen and bone marrow of day 21 Dox treated mice had significantly lower proliferation compared to controls (Fig. 5D). To test if this decrease was due to a change in differentiation of the leukemia cells, we examined markers of mature myeloid cells, Mac-1 and Gr-1. In the spleen of Dox treated mice, we observed an increase in Mac1^+^, GR-1^+^ leukemia cells, consistent with differentiation. In contrast, there was no difference in the presence of Mac1 or GR-1 on the bone marrow leukemia cells (Fig. 5E). To test whether bone marrow *CM* KD cells were instead enriched for leukemia stem cell (LSC) markers, we examined expression of KIT, CD59a, IL1RL1, and CD123. Work by us and others previously showed that KIT, CD59a and IL1RL1 are enriched on the LSC population driven by *CM* expression, and CD123 is an established LSC marker in human AML (25, 32, 33). We observed a small yet statistically significant increase in CD123 and decrease in IL1RL1 in the leukemia cells in the bone marrow of Dox treated mice (Fig. 5F). These results indicate that the decreased proliferation observed in the bone marrow *CM* KD cells is not caused by differentiation to a post-mitotic mature myeloid cell or enrichment for quiescent LSCs.

### KD of *CM* is associated with distinct changes in gene expression in the spleen and bone marrow

To investigate the transcriptomic differences associated with the survival of *CM* KD leukemic cells in the bone marrow, but not the spleen, we performed bulk RNA sequencing (RNA-Seq) on GFP^+^ leukemia cells from both populations on day 7 of control or Dox treated mice. We identified a total of 348 differentially expressed genes (DEGs); (221 upregulated and 127 downregulated) in leukemia cells from the spleen of Dox treated animals, as compared to control. In leukemia cells from the bone marrow, we identified a total of 4650 DEGs (2328 upregulated and 2322 downregulated) in cells from Dox treated mice, as compared with control. (Supplementary Table 2). Hierarchical clustering shows the bone marrow replicates from control and Dox treated mice have similar DEG patterns (Supplementary Fig. S4A). While spleen control replicates cluster together, one Dox treated sample clustered more closely with the control spleen samples, suggesting an intermediate level of transcriptional changes, perhaps related to inefficient KD of the fusion protein (Supplementary Fig. S4B, C). In the bone marrow, several known targets of the CM fusion protein, including *Cebpe, Cebpd, Runx1 and Il1rl1*, are downregulated with *CM* KD. In addition, genes associated with an abnormal myeloid progenitor (AMP) phenotype, including *Rhag, Cldn13, Add2, Gata2* were found to be upregulated (32). Overall, these changes in genes expression are consistent with loss of CM activity but imply that *CM* KD cells retain transcriptional changes associated with leukemic blasts (Fig. 6A).

**Figure 6:**
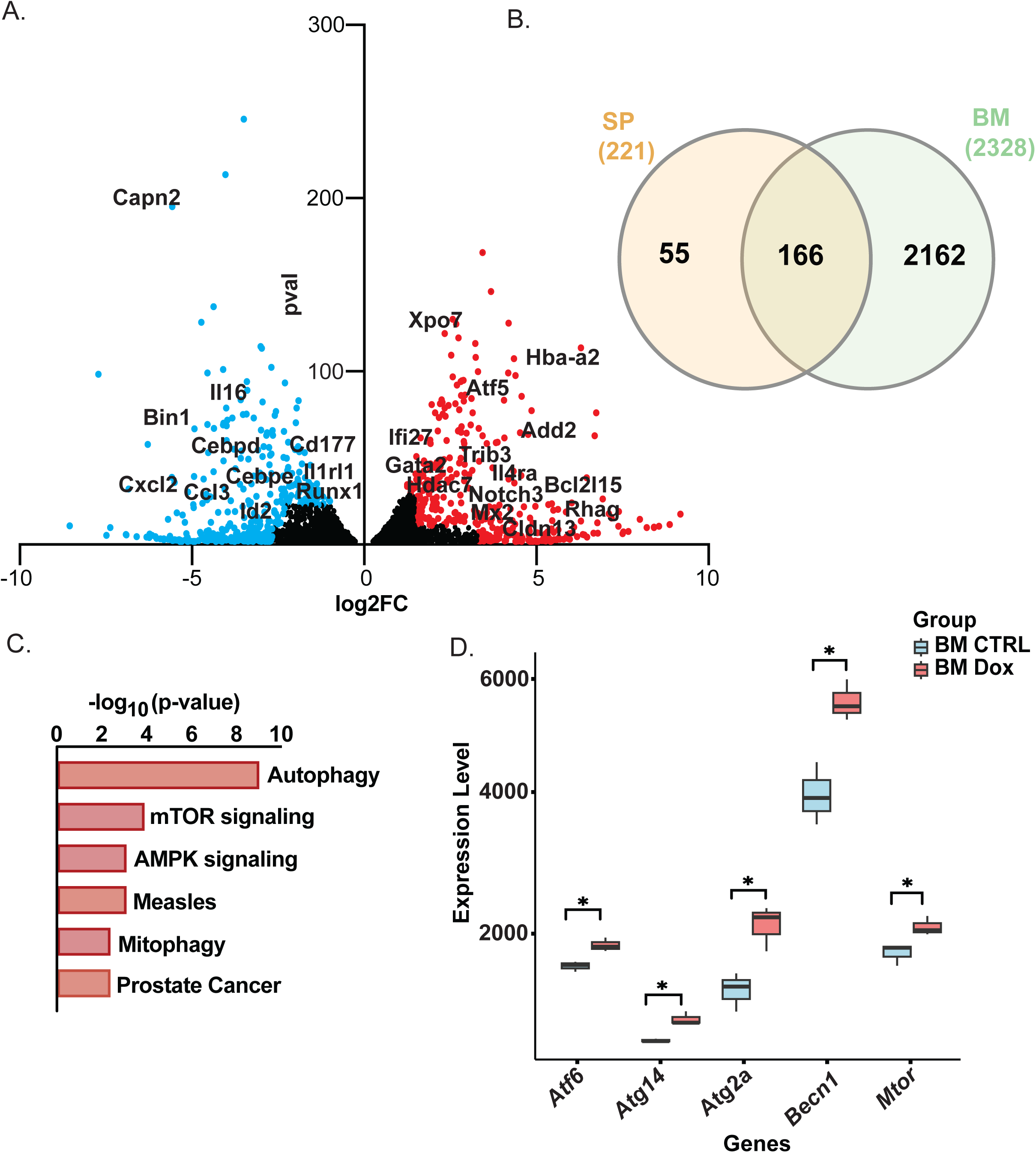
Increased expression of autophagy related genes in *CM* knockdown leukemia cells from the bone marrow. **A.** Volcano plot showing distribution of the statistically significant differentially expressed genes (DEGs) in the GFP^+^ sorted leukemia cells from the BM of mice treated for 7 days with CTRL or Dox. A few of the key genes with a log_2_ fold change ≥ 0.5 and adjusted p value ≤ 0.05 are highlighted. **B**. Venn diagram showing overlap of significantly upregulated DEGs in *CM* KD leukemia cells from the SP and BM, as compared to CTRL. **C.** Pathway analysis of upregulated DEGs in *CM* KD cells from the BM showing the top KEGG pathways affected. **D.** Box plot depicting relative expression of indicated autophagy associated genes in control and *CM* KD leukemia cells from the BM. N = 3 biological replicates, * = padj ≤ 0.05 as compared to CTRL.

To identify biological functions that promote survival of bone marrow *CM* KD cells, we compared DEGs in spleen and bone marrow leukemia cells and performed pathway analysis on only those genes altered in the bone marrow. Of the 2162 unique DEGs upregulated in bone marrow leukemia cells we identified autophagy, a known promoter of AML survival, as well as mTOR signaling, AMPK signaling, and mitophagy which have been implicated in promoting autophagy, survival, and stress adaptation in AML (Fig 6C) (34–38). Analysis of individual autophagy related genes showed significantly increased expression of *Atf6*, *Atg14*, *Atg2A*, *Becn1* and *mTor* (Fig 6D). These results imply a role for autophagy in the survival of *CM* KD cells in the bone marrow. Pathway analysis of leukemia cells from the spleen indicated that the top deregulated pathway with KD of *CM* is apoptosis, consistent with the Annexin V staining and western blot analysis in Figure 5 (Supplementary Fig. 4C).

To determine if *CM* KD induces distinct gene expression changes in different subpopulations of leukemia cells, we performed single-cell RNA sequencing (scRNA-Seq). We harvested cells from the bone marrow and spleen of control and dox treated mice at day 7 of treatment. Single cell suspensions were lineage depleted and analyzed by flow cytometry to confirm enrichment of leukemia cells. All samples showed >60% leukemia cells, with the bone marrow approaching 80% in both control and Dox treated samples (Supplementary Fig. 5A). Single-cell transcript libraries were prepared and sequenced.

Unbiased clustering analysis revealed multiple subpopulations of leukemia cells in both the spleen and bone marrow. Interestingly, *CM* KD affected the relative proportion of these clusters, with the number of cells increased in some clusters while others decreased in samples from Dox treated mice (Fig 7A, B). In the spleen, Clusters 2 and 4 appeared to be the most sensitive to *CM* KD (Supplementary Fig. 5B,C). In the bone marrow, Clusters 3, 4, 5, 6, 9 and 10 showed marked reduction with *CM* KD while the number of cells in Clusters 0, 1, 2, 7 and 8 increased, indicating these clusters don’t require *CM* expression for survival (Fig. 7B). This analysis shows that there is dramatic cellular heterogeneity among the leukemia cells in both the bone marrow and spleen, with different subpopulations differentially dependent on *CM* expression.

**Figure 7:**
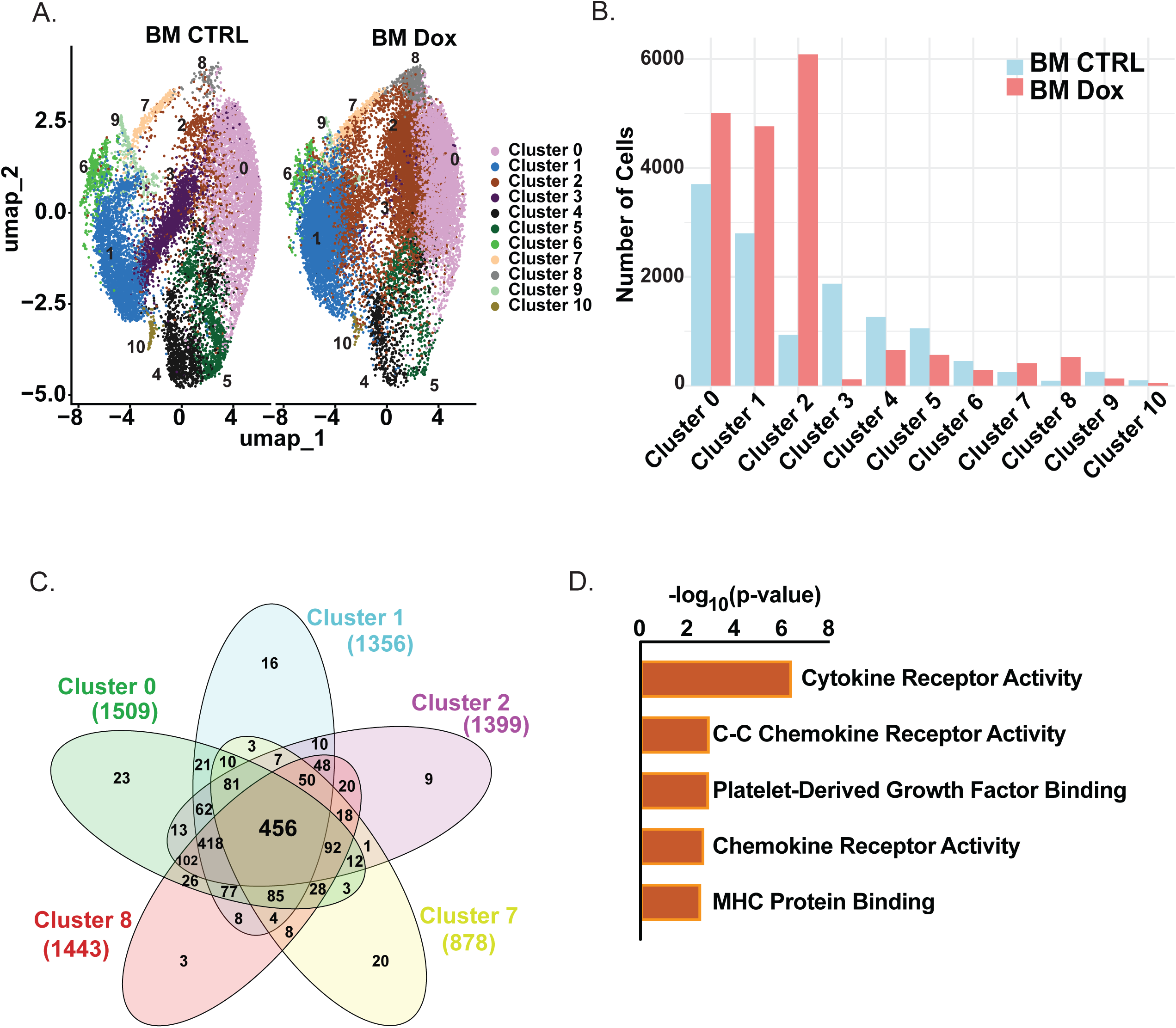
Single cell transcriptomics reveals distinct *CM*-dependent and *CM-*independent subpopulations in the bone marrow. **A.** Umap projection of leukemia cells from the BM of CTRL or Dox treated mice. Each dot represents a single cell and is colored according to the assigned transcriptional cluster. **B.** Bar graph of the number of cells in each cluster in BM samples from CTRL or Dox treated mice **C.** Venn diagram representing the shared genes expressed by cells in the *CM* independent clusters in BM samples from CTRL mice with 456 genes common among the five groups. **D.** Top 5 molecular functions associated with the 456 shared genes from C.

To test if there are inherent transcriptional programs promoting the survival of the *CM*-independent cells in the bone marrow, we compared the transcripts expressed in the clusters that don’t require CM expression. We identified 456 genes expressed in all *CM*-independent clusters (Fig 7C). Among these, several genes including *Ccl5, Ccr1, S100a4, CD74, CD27, Ccnd2* and *Cdkn1a* are all associated with leukemia survival and proliferation (39–43). Consistent with this finding, pathway analysis indicates that these genes are significantly enriched for molecular functions involving cytokine and chemokine activity (Fig. 7D). In contrast, the common genes expressed in *CM-*dependent clusters were associated with non-immune molecular functions (Supplementary Fig. 5D). Collectively these results imply that cytokine and chemokine signaling within the bone marrow microenvironment may be a critical mechanism promoting survival of *CM*-independent leukemia cells.

### *CM* KD leukemia cells can re-establish disease

The persistence of a *CM* KD leukemic population in the bone marrow raises the possibility that these cells may be able to cause relapsed disease. To test this possibility, we maintained mice transplanted with *shMYH11* transduced leukemia cells on Dox or control drinking water until they showed signs of lethal disease. 11 out of 18 Dox treated mice showed re-emergence of GFP^+^ cells in the blood (Fig. 8A). To determine if these cells expressed *CM*, we sorted for GFP^+^ cells and performed qRT-PCR and western blot (Fig. 8 B-D). We found that GFP^+^ relapse cells had reduced *CM* at both the mRNA and protein levels, as compared to parental leukemia cells from control treated mice. In addition, we found that the immunophenotype of relapsed *CM* KD leukemia cells was similar to parental leukemia cells from control mice (Supplementary Fig. S6A and B). To understand the molecular mechanism allowing *CM* KD cells to re-emerge into the blood, we performed whole transcriptome sequencing on relapse and parental leukemia cells. The leukemic cells from relapsed mice were harvested from the spleen and compared with either spleen or bone marrow from control mice. Principal component analysis (PCA) showed that relapsed samples did not cluster with parental leukemia cells harvested from either the bone marrow or spleen indicating that relapse is accompanied by widespread alterations in transcription (Fig. 8E). In addition, only 3 of the 4 relapsed samples clustered together, implying that there may be multiple transcriptional programs that promote re-emergence of *CM* KD cells from the bone marrow. Comparing the transcriptome of relapsed samples to control, we observed a total of 930 DEGs (359 upregulated and 571 downregulated) in relapsed samples compared to parental leukemia cells (Supplementary Table 3). Further, pathway analysis indicates that the DEGs are associated with multiple pathways implicated in leukemia survival and relapse, including interferon regulator factor (IRF) signaling (Fig. 8F). Specifically, *Irf1*, *Irf7*, and *Irf9* are decreased in relapse samples compared to controls (Fig. 8G). Notably, loss of these genes is associated with relapse in AML patients (46–48). This implies that suppression of IRF signaling allows *CM* KD leukemia cells to survive outside the bone marrow.

**Figure 8:**
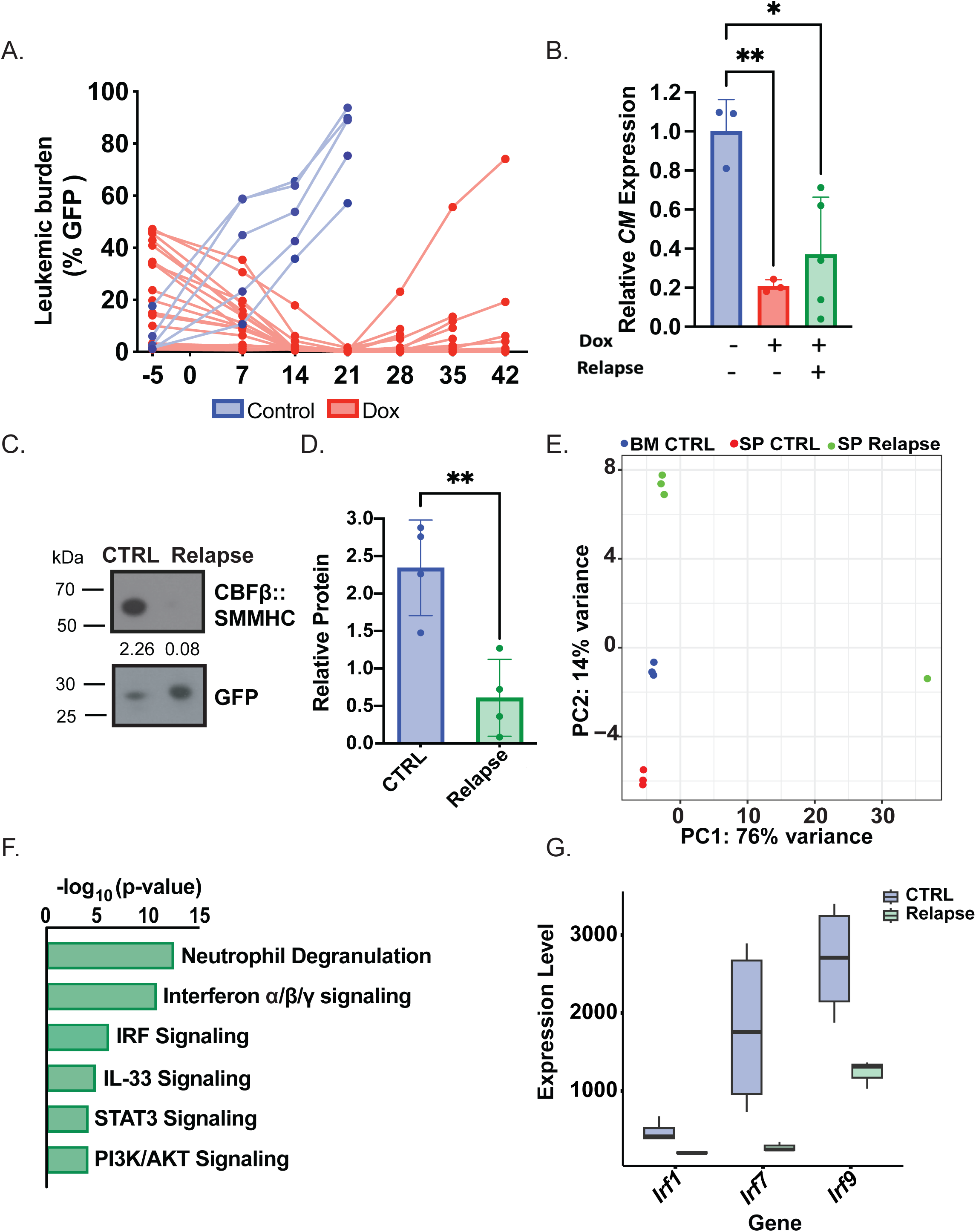
The surviving *CM* knockdown cells in the bone marrow can give rise to relapse. **A.** Line graph showing the % GFP^+^, *shMYH11* transduced cells in PB of mice treated for up to 42 days with either CTRL or Dox**. B.** Bar graph of relative *CM* expression in GFP^+^ sorted leukemia cells from Dox treated mice after relapse. **C.** Representative western blot and **D.** Bar graph of relative CBFβ::SMMHC levels from sorted GFP^+^ cells harvested from CTRL or Dox treated mice. Quantification to GFP. **E.** Principal Component Analysis (PCA) plots of whole transcriptome sequencing data from GFP sorted relapse or CTRL leukemic cells harvested from the SP or BM. **F.** Pathway analysis of the differentially expressed genes in relapse samples compared to the combined SP and BM CTRL leukemia samples. **G.** Bar graph depicting the *Irf* genes deregulated in Dox treated mice after relapse. N ≥ 3 biological replicates, * = p ≤ 0.05 as compared to CTRL.

Collectively, this study indicates that while *CM* is required for the survival of most frank leukemia cells, a population of leukemia cells in the bone marrow do not strictly require *CM* activity and can re-establish fatal disease.

## Discussion

Given that CBFβ::SMMHC is expressed in all leukemia cells of Inv(16) AML patients, but not in healthy cells it is a tempting target for drug development. However, most studies have focused on the role of the fusion protein during leukemia initiation and less is known about its role in frank leukemia cells in the context of cooperating mutations. Previous work in the Inv(16) immortalized cell line, ME-1, reported that decreased expression of *CM* caused changes in gene expression consistent with differentiation, but did not cause any changes in viability at the time point analyzed, raising the possibility that continued *CM* expression may not be strictly required for leukemia maintenance (27). To our knowledge, ours is the first study to test whether *CM* activity is required for leukemia cell survival using non-immortalized leukemia cells. We found that KD of *CM* induced apoptosis and decreased colony forming ability *in vitro*. *CM* KD *in vivo* caused a near complete elimination of leukemia cells in the blood and spleen. Surprisingly, the leukemic burden in the bone marrow did not decrease below 20-30% despite almost complete loss of the CBFβ::SMMHC protein, implying that the bone marrow microenvironment can sustain leukemia cells despite the loss of *CM* expression.

Our observation that bone marrow *CM* KD cells, but not spleen, had decreased apoptosis as compared to controls suggests tissue specific effects on the survival of leukemic cells *in viv*o. Thus, to identify potential factors promoting the survival of bone marrow *CM* KD cells, we analyzed the anti-apoptotic proteins BCL-2 and MCL-1 (44, 45). However, we observed that both control and *CM* KD leukemic cells had robust expression of MCL-1 but nearly undetectable levels of BCL-2, indicating that differential expression of anti-apoptotic proteins does not explain their survival. Rather, our bulk RNA-seq indicates that the upregulation of autophagy may contribute to the survival of bone marrow *CM* KD cells. These findings are consistent with recent work showing a role for autophagy in leukemia progression and that overexpression of the autophagy genes *BECLIN1* and *ATG* are associated with shorter overall survival in AML patients (46, 47).

To understand the cellular heterogeneity in *CM* sufficient and KD cells, we performed scRNA-Seq. We found that there are clusters of leukemia cells with distinct transcriptional profiles and different requirements for *CM* expression. Some clusters had decreased cellularity while others were increased, implying the presence of *CM*-dependent and *CM*-independent leukemic populations in the bone marrow. This is consistent with our *in vivo* results where we observed an initial decrease in bone marrow leukemic burden of dox treated mice, but eventual stabilization of the leukemia population at 20-30%. Interestingly, the population of *CM*-independent leukemia cells did not continue to increase over time, implying that there is a maximum number of *CM*-independent leukemia cells that the bone marrow can support. Of note, the *CM*-independent clusters in the bone marrow shared a transcriptional profile associated with cytokine and chemokine activity, which is consistent with a model in which the bone marrow microenvironment provides a limited amount of survival factors, keeping the *CM-*independent population from expanding indefinitely in the bone marrow.

Our results concerning the role of CBFβ::SMMHC during leukemia maintenance are strikingly similar to what has been demonstrated with the BCR::ABL fusion protein in CML. Pharmacological inhibitors of BCR::ABL, such as imatinib, are remarkably successful at inducing long-term, stable cytogenetic remission in CML (48, 49). However, these kinase inhibitors generally do not eliminate the LSC population in the bone marrow. Further, transgenic models have demonstrated that with near complete knockdown of *BCR::ABL* expression, LSCs remain in the bone marrow and are able to re-establish disease when BCR::ABL activity is restored (50, 51). In our model, we also observed a similar population of LSC-like cells that can survive in the bone marrow and re-establish disease, despite near complete loss of the CBFβ::SMMHC protein. In our case, relapse did not require re-expression of the fusion protein. However, given the period of latency before relapse, it is likely that additional mutations developed to compensate for the lack of CBFβ::SMMHC activity.

Collectively, this study demonstrates that *CM* expression continues to be required after leukemic transformation and supports the development of inhibitors targeting the fusion protein for the treatment of Inv(16) AML. Our data indicates such inhibitors would be effective at managing leukemic burden and, when used in combination with other therapeutics, have the potential to cure the disease.

## Acknowledgments

The authors would like to thank Paul Liu, M.D, Ph.D (NHGRI/NIH) for his support and encouragement of this project, as well as the members of the UNMC flow cytometry and Bioinformatics and Systems Biology Core Facilities, and the Nebraska Center for Molecular Target Discovery and Development for their expertise.

## Author Contributions

S.P. designed and performed experiments, analyzed data and wrote the paper. Y.W. designed and performed experiments and analyzed data. M.B., C.L., A.D., and S.A.S. performed experiments. R.D. and P.W analyzed single cell and bulk RNA-Seq data, respectively. C.R., L.A., and L.G. generated, bred, and mutagenized *Cbfb^+/floxMYH11^* knock-in mice. S.A.S., K.J.H. and R.K.H designed experiments, analyzed data and wrote the paper.

## Disclosure of Conflicts of Interest

The authors have no conflicts of interest to disclose.

## Notes

This work was supported by R00 CA148963 and R01 CA244900 to R.K.H and T32 CA009467 to support C.L. C.R., L.A., and L.G. were supported by the Intramural Research Programs of the National Human Genome Research Institute, National Institutes of Health. Core facilities used for this project are supported by a state fund from the Nebraska Research Initiative, the Fred and Pamela Buffett Cancer Center’s National Cancer Institute Cancer Support Grant (P30 CA036727), and the Nebraska Center for Molecular Target Discovery and Development COBRE (P20 GM121316).

### Competing Interest Statement

The authors have declared no competing interest.

### Summary of Updates

This version of the manuscript has been revised to include additional findings.

