## Supplementary Materials for "Tissue Specific Requirement for the Inv(16) Oncogene *CBFB::MYH11* in Acute Myeloid Leukemia"

#### Supplemental Methods

##### Whole Transcriptome sequencing

The trimmed fastq files were processed by utilizing STAR and RSEM (1) for quantification at gene level (2). Reads were mapped to GRCM38 mouse reference genome. The raw counts were used for differential expressed gene (DEGs) analysis by DESeq2 with a padj ≤ 0.05 and gene results were sorted by $\log_2$FC (3). Heat maps were plotted by gplots v3.1.3.1 package in R 4.4.0 for all significant genes with padj ≤ 0.05 in all samples. Functional categorization was performed using EnrichR with padj ≤ 0.05 (4). The tow bar plots were generated using plotted ggplot2 3.5.2 and expression levels are normalized to DESeq2 expression.

##### Single cell RNA Sequencing

Library preparations were made as previously described (5) with following modifications to improve product yield. For the reactions 300 pg/ul was made. After initial centrifugation samples were incubated at 55 C for 3 minutes. All of the steps until adding Neutralization buffer were performed on ice. Quality of whole transcriptome libraries were checked using a D1000 TapeStation (Agilent) and sequenced. The replicates were pooled for the sequencing. Seurat library from R Bioconductor (6) has been used to perform the filtering and processing of the samples. Clusters were identified in an unbiased manner.

### Supplementary Table 1

| Antibody | Manufacturer, City, Town, Country |
|---|---|
| MYH11 | Origene, Rockville, MD, USA |
| GAPDH | Ambion, Austin, TX, USA |
| β-ACTIN | Santa Cruz Biotechnology, Dallas, TX, USA |
| MCL-1 | Cell Signaling Technology (CST), Danvers, MA, USA |
| BCL-2 | CST |
| GFP | Invitrogen, Waltham, MA, USA |
| Anti-Mouse | Vector Laboratories, Newark, CA, USA |
| Anti-Rabbit | Vector Laboratories, Newark, CA, USA |

#### Supplemental Figure 1

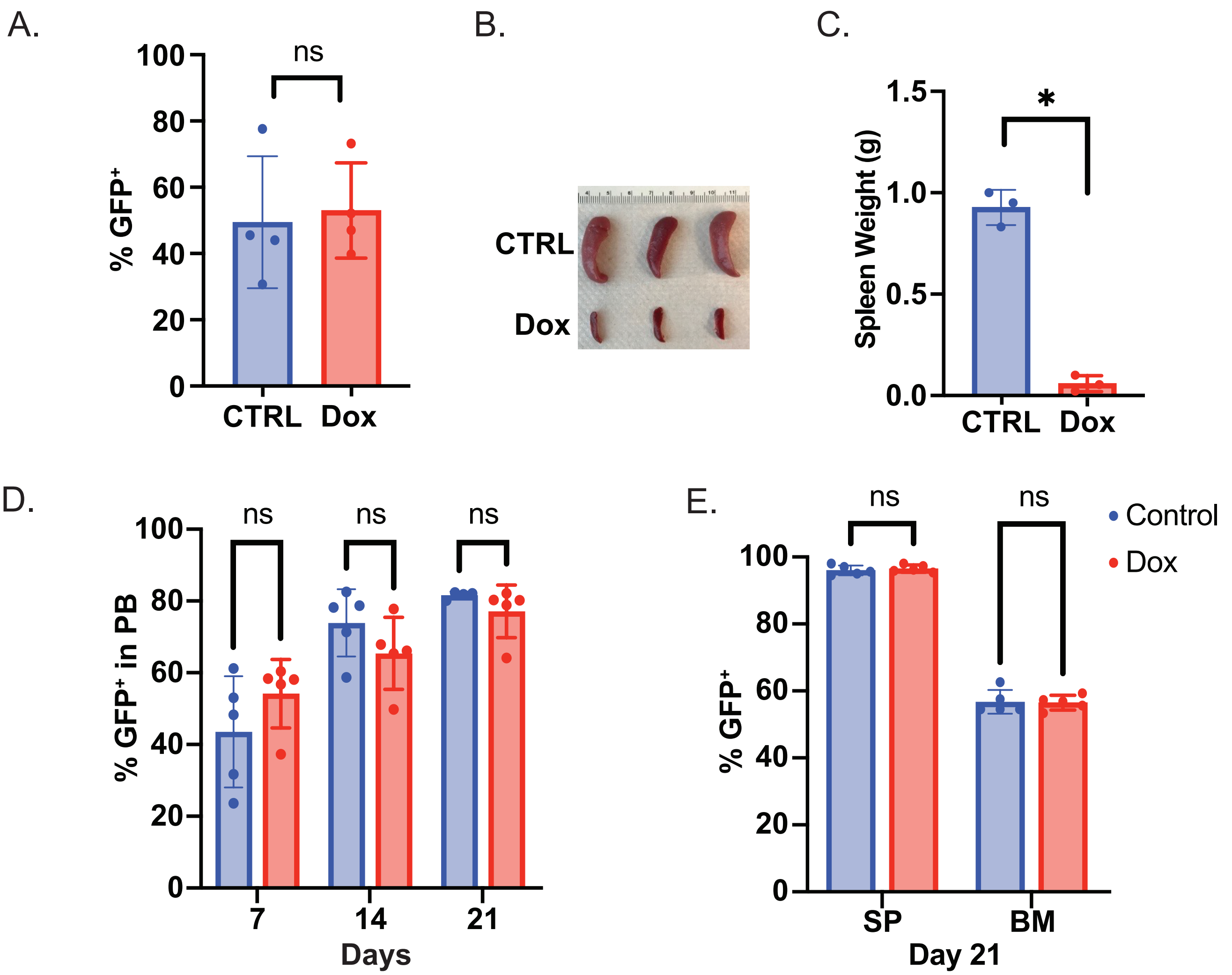


**Supplementary Figure S1: Knockdown of CM in vivo decreases splenomegaly. A.** Bar graph showing the percentage (%) of GFP+ cells in the peripheral blood (PB) before treatment with control (CTRL) or doxycycline (Dox) for representative mice. **B**. Representative photograph of spleens harvested from mice treated with CTRL or Dox for 21 days. **C.** Bar graph of spleen weights from mice treated as in B. **D.** Bar graph showing the % GFP+ cells in the PB after treatment with CTRL or Dox at days 7, 14 and 21 in the non-shMYH11 GFP clone. **E.** Bar graph showing the % GFP+ cells in the spleen (SP) and bone marrow (BM) of the representative mice in D. sacrificed at day 21 of treatment. N ≥ 3, ns = non-significant, * = $p \leq 0.0001$ as compared to CTRL.

#### Supplemental Figure 2

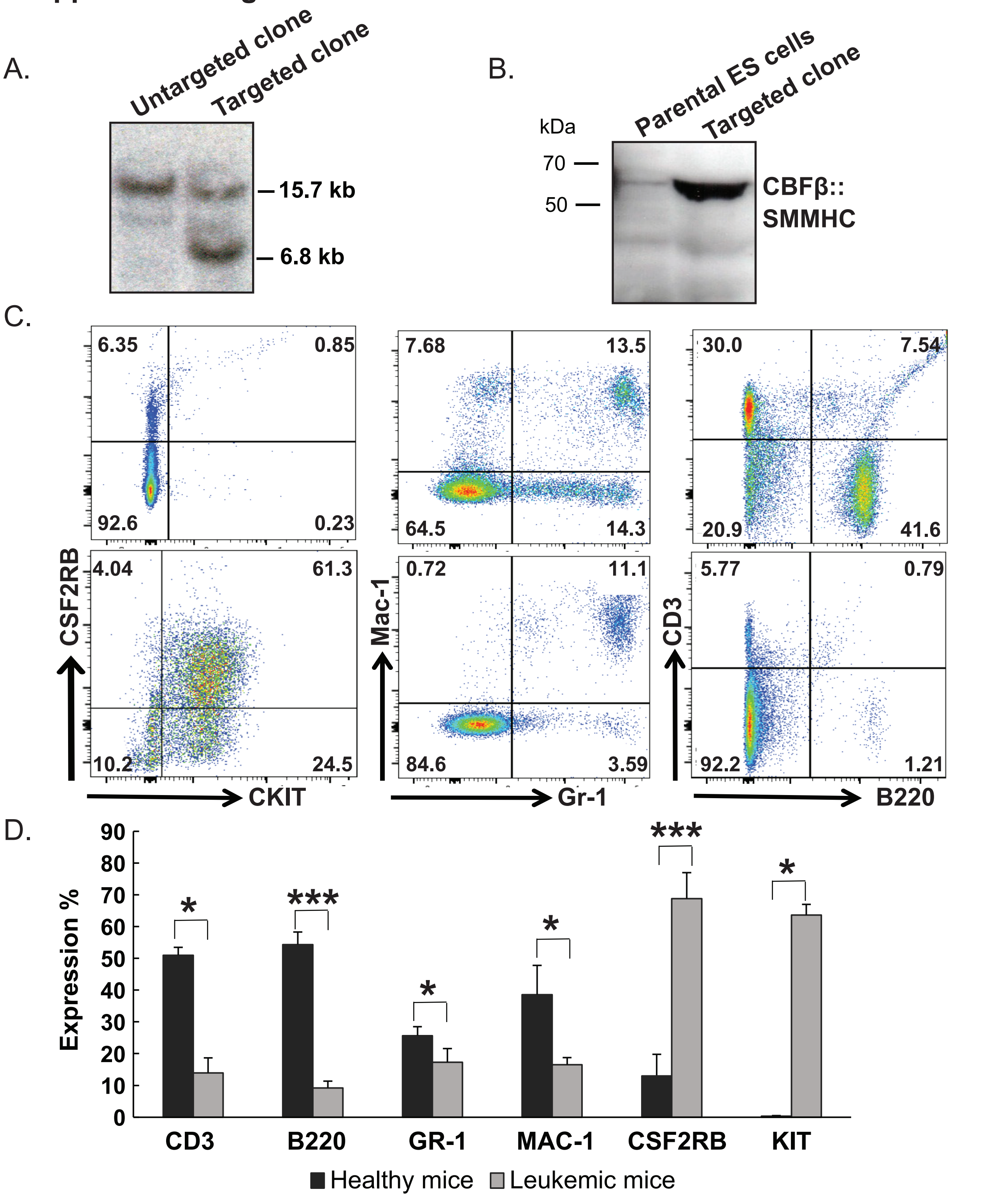


**Supplementary Figure S2: Mice with the f*loxCbfb::MYH11* allele develop myeloid leukemia. A.** Representative northern and **B.** western blots from targeted and untargeted embryonic stem cell clones. **C.** Representative cell surface staining with the indicated antibodies of PB from healthy or leukemic recipient mice transplanted with leukemic cells from a *Cbfb*$^{+/flMYH11}$ chimeric mouse. **D.** Bar graph indicating the percentages of blood cells expressing CD3, B220, MAC-1, GR-1, CSF2RB, and KIT from mice transplanted with CbfbfloxMYH11 leukemic cells. N = 3. * = p ≤ 0.05; ** = p ≤ 0.01; *** = p ≤ 0.001 as compared to CTRL.

#### Supplemental Figure 3

A.

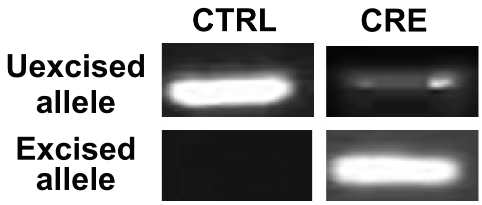


B.

| | Total # of colonies analyzed | #of colonies with *Cbfb::MYH11* deletion | Frequency of colonies with *Cbfb::MYH11* deletion |
|---|---|---|---|
| Chimera #1 | 14 | 2 | 14 % |
| Chimera #2 | 5 | 1 | 20 % |
| Chimera #3 | 20 | 1 | 5 % |

**Supplementary Figure S3: Leukemia cells lacking *CM* can form colonies *in vitro*. A.** Representative PCR of the excised and unexcised $Cbfb^{floxMYH11}$ allele from a single colony grown from cells transduced with either control or Cre expressing virus. **B.** Table showing the results of single colony PCR grown from Cre transduced leukemia cells from 3 different $Cbfb^{floxMYH11}$ chimeric mice.

#### Supplemental Figure 4

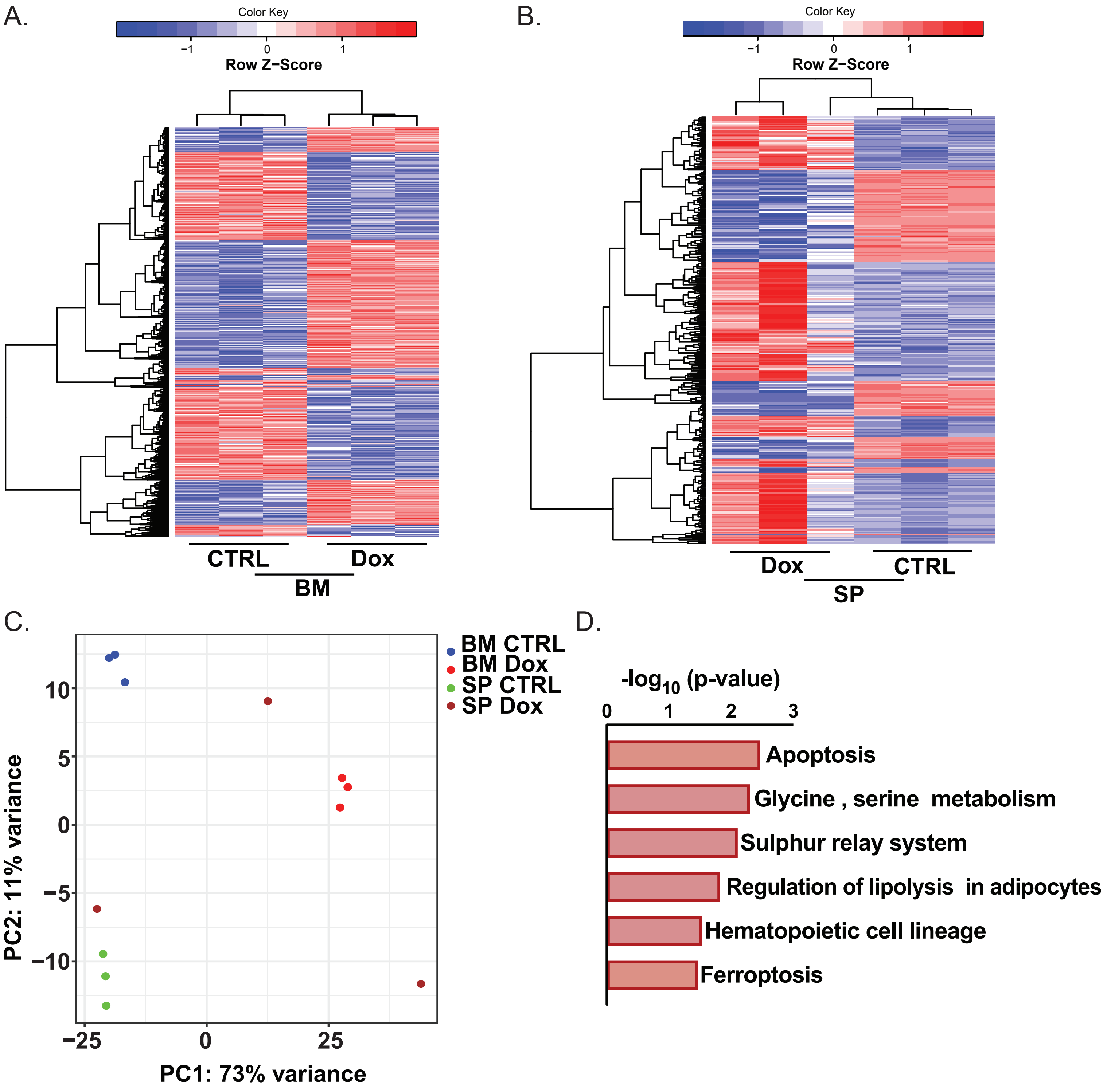


**Supplementary Figure S4: CM KD cells from the spleen show upregulation of genes associated with apoptosis. A.** Heat map of the significantly differentially expressed genes (DEGs) in the GFP+ sorted leukemia cells from the BM and **B.** SP of mice treated for 7 days with CTRL or Dox. The rows represent the genes and column represents the biological replicates from both the groups. **C.** Principal Component Analysis (PCA) plot of gene expression profiles of the cells from A. **D.** Pathways enrichment analysis of DEGs in CM KD cells from the spleen showing the top pathways affected. N = 3 biological replicates, padj ≤ 0.05 as compared to CTRL.

#### Supplemental Figure 5

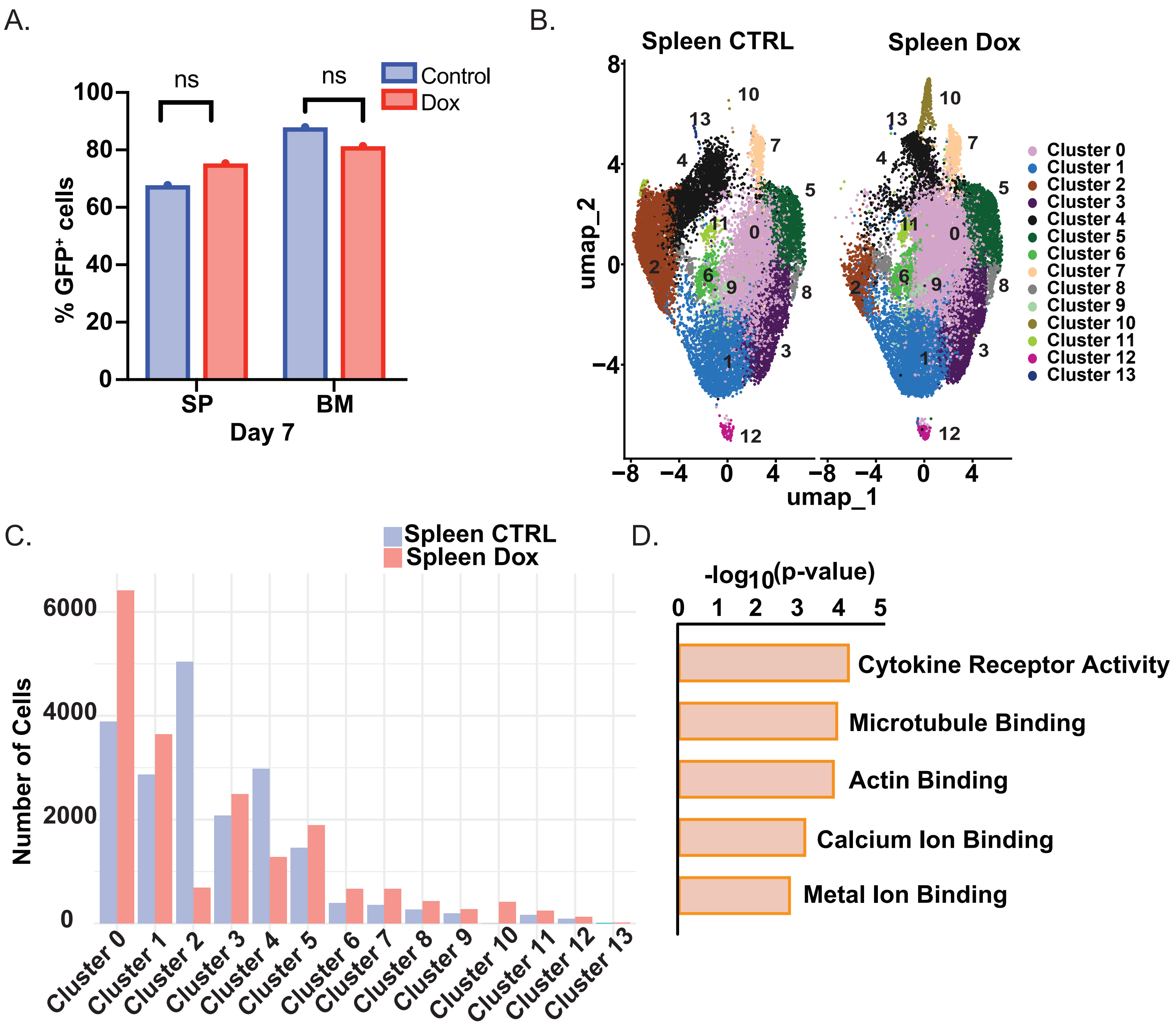


**Supplementary Figure S5: Single-cell transcriptomics reveals distinct CM-dependent and CM-independent subpopulations in the spleen. A.** Bar graphs showing the % GFP+ leukemia cells from lineage depleted SP or BM samples from CTRL or Dox treated mice after 7 days. **B.** Umap projection of leukemia cells from the SP of CTRL or Dox treated mice. Each dot represents a single cell and is colored according to the assigned transcriptional cluster. **C.** Bar graph of the number of cells in each cluster from B. **D.** Top 5 molecular functions associated with the 1752 genes shared by cells in the *CM*-dependent samples from CTRL mice.

#### Supplemental Figure 6

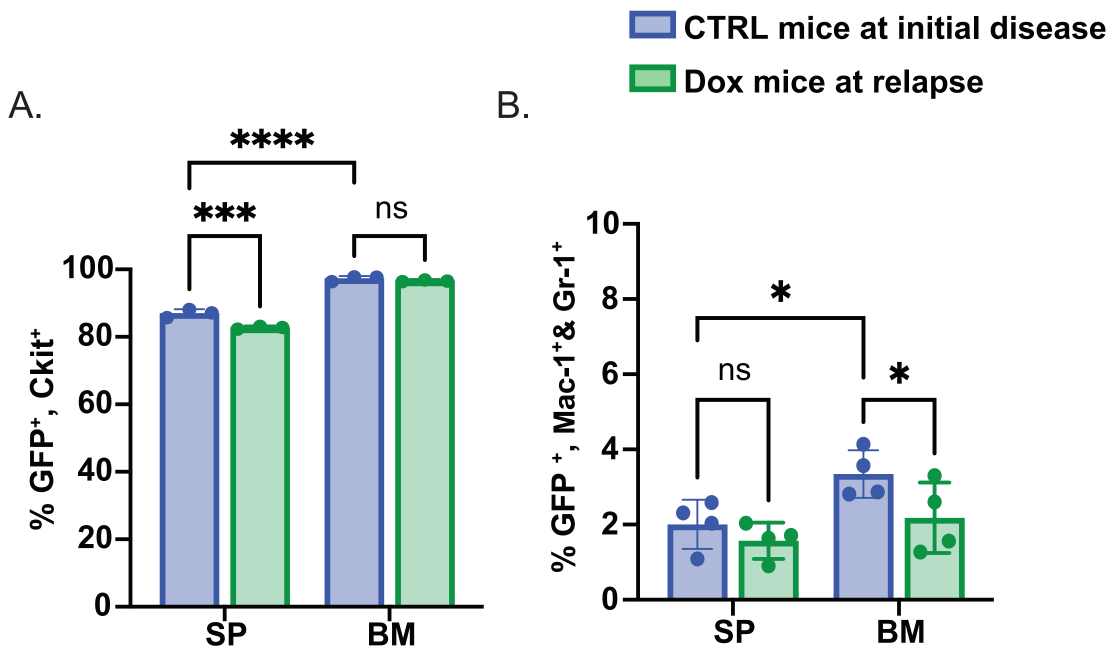


**Supplementary Figure S6: *CM* KD leukemia cells at relapse have a similar immunophenotype to control leukemia cells. A.** Bar graph showing the % GFP+ cells expressing Ckit in the SP and BM of CTRL mice during initial disease and Dox mice at relapse. **B.** Bar graph showing % GFP+ cells expressing the mature myeloid markers Mac-1 and Gr-1 harvested from the SP and BM of mice treated as in A. N ≥ 3 biological and technical replicates, * = p≤0.05 as compared to CTRL.
